# Abstinence and Extinction Drive Opposing Changes in Striatal Activity and Dopamine Signaling During Alcohol Relapse

**DOI:** 10.1101/2025.06.02.657507

**Authors:** Xueyi Xie, Ruifeng Chen, Xuehua Wang, Jun Wang

## Abstract

Relapse remains a major obstacle in the treatment of alcohol use disorder, often driven in part by enduring neuroadaptations. However, how different treatment strategies—such as abstinence versus extinction training—modulate the underlying neural circuits and synaptic mechanisms that shape relapse vulnerability remains poorly understood. In this study, we demonstrate that abstinence and extinction distinctly influence dorsomedial striatal (DMS) direct-pathway medium spiny neuron (dMSN) activity and dopamine signaling during cue-induced reinstatement of alcohol seeking. Using in vivo fiber photometry in D1-Cre rats expressing calcium or dopamine sensors, we found that abstinence enhanced dMSN calcium responses and dopamine release during reinstatement, whereas extinction normalized these neural signals and suppressed relapse-like behavior. Furthermore, bidirectional optogenetic modulation of medial prefrontal cortex (mPFC)–to–dMSN synapses revealed a causal role for corticostriatal plasticity in determining relapse propensity. Inducing long-term depression (LTD) in the abstinent state attenuated reinstatement, while inducing long-term potentiation (LTP) after extinction training reinstated alcohol seeking. Together, these findings identify distinct neural adaptations shaped by abstinence versus extinction and highlight corticostriatal plasticity as a potential target for relapse prevention.

**Highlight:**

1. Abstinence enhances striatal dMSN activity and dopamine signaling during cued relapse.
2. Extinction training normalizes dMSN dynamics and reduces dopamine release during cued relapse.
3. Optogenetic mPFC-to-dMSN long-term depression after abstinence reduces relapse.
4. Optogenetic mPFC-to-dMSN long-term potentiation after extinction invigorates relapse.

## Introduction

Relapse remains a major challenge in the treatment of alcohol use disorder (AUD).^1,2^ Although sustained abstinence is the optimal outcome for individuals with AUD,^3^ forced cessation of alcohol use often triggers progressively greater craving over time, increasing vulnerability to relapse.^4–6^ Behavioral interventions such as extinction training have shown promise in reducing relapse vulnerability.^7,8^ In clinical populations, extinction-based approaches can attenuate striatal responses to alcohol-related cues, while in animal models, extinction training reliably suppresses relapse-like behavior.^9–13^ Mechanistically, extinction of cocaine seeking has been shown to reduce glutamatergic synaptic transmission onto ventral striatal dMSNs, in contrast to abstinence, which enhances excitatory input to this population.^14^ Extinction of heroin seeking also dampens thalamostriatal plasticity in ventral striatal dMSNs.^15^ However, despite these insights, the precise neural mechanisms by which extinction alters striatal circuit function to suppress alcohol relapse remain poorly understood, limiting its broader translational application.

The striatum, particularly its dorsomedial region (dorsomedial striatum, DMS), plays a critical role in encoding goal-directed behaviors and driving relapse to alcohol seeking.^16–19^ The principal neurons within the striatum are medium spiny neurons (MSNs), which are classically divided into two major subtypes based on their dopamine receptor expression and projection targets.^20,21^ Direct-pathway MSNs (dMSNs) express dopamine D1 receptors and project directly to the substantia nigra pars reticulata; their activation facilitates positive reinforcement and promotes alcohol consumption.^21–23^ In contrast, indirect-pathway MSNs (iMSNs) express dopamine D2 receptors and project to the external segment of the globus pallidus; their activation is associated with aversive outcomes and suppression of alcohol drinking.^22–25^

All addictive substances, including alcohol, trigger striatal dopamine release,^26^ which preferentially activates dMSNs and strengthens their corticostriatal synaptic connections.^18,27–29^ Notably, optogenetic inhibition of dorsostriatal dMSNs has been shown to suppress operant alcohol seeking and reduce relapse.^30^ Moreover, alcohol-induced synaptic plasticity between the medial prefrontal cortex (mPFC) and dMSNs plays a bidirectional role in modulating operant alcohol-seeking behaviors.^31–33^ Despite these advances, the precise neural dynamics of dMSN activity and dopamine release during operant self-administration (OSA) of alcohol remain unclear. Furthermore, how abstinence or extinction experiences influence these dynamics has yet to be fully elucidated.

In this study, we employed genetically encoded sensors to monitor real-time activity of DMS dMSNs and dopamine release dynamics in freely moving rats during operant self-administration (OSA) of alcohol. We further extended these measurements to relapse conditions following either forced abstinence or extinction training to determine how these experiences shape striatal signaling. Lastly, we utilized dual-channel optogenetics to bidirectionally manipulate corticostriatal plasticity after abstinence or extinction, and tested its causal role in regulating relapse vulnerability. Together, these experiments aim to uncover circuit- and synapse-level mechanisms underlying alcohol relapse and provide insights into targeted interventions that may prevent it.

## Results

### DMS dMSN activity and dopamine release peak after lever press but before alcohol consumption during operant self-administration

Striatal dMSNs are preferentially activated during substance use, likely via enhanced dopamine release.^28,34,35^ However, the precise temporal dynamics of dMSN activity and dopamine release during alcohol OSA remain unclear. To address this, we first examined when and how DMS dMSNs are activated, and when dopamine is released to the DMS during alcohol OSA. This paradigm closely models human patterns of alcohol use and relapse.^36,37^

In our paradigm, rats were trained to press an active lever to receive alcohol, which was delivered into a central magazine (Fig. 1A). A discrete cue light within the delivery port accompanied each delivery and was later used to induce relapse. To monitor neural dynamics, we unilaterally expressed AAV-FLEX-jGCaMP7f in DMS dMSNs and AAV-GRAB_DA2m_ in the contralateral DMS for dual-site fiber photometry recordings in D1-Cre rats (Fig. 1B-1D).

**Figure 1.**
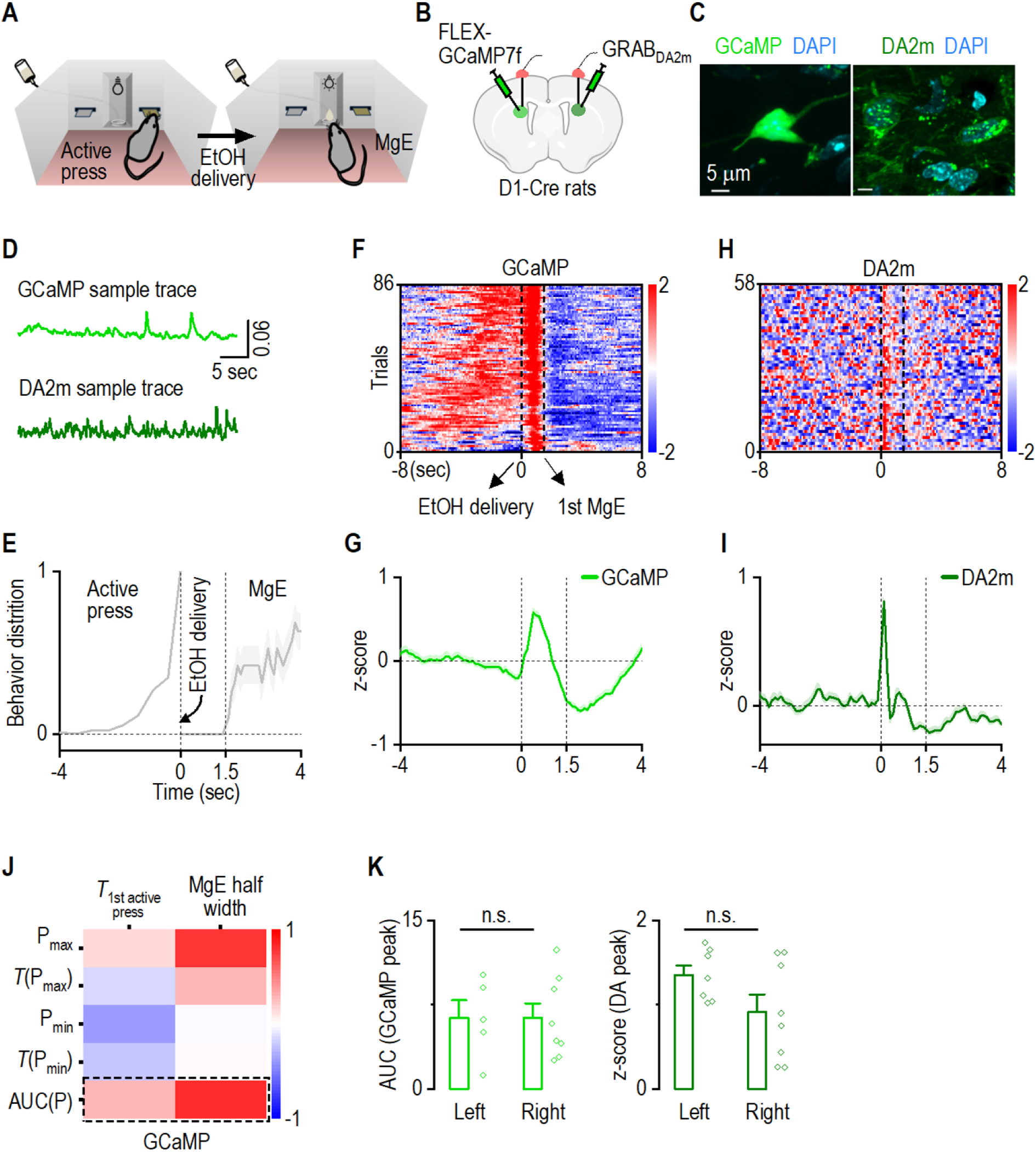
DMS dMSN calcium activity and dopamine release peak after active presses but before magazine entries during operant self-administration of alcohol. *A*, Schematic of the operant self-administration (OSA) paradigm for alcohol (EtOH). Rats were trained to press an active lever to receive 0.1 mL of 20% (v/v) alcohol, delivered alongside a cue light above the port. Rats then entered the magazine (magazine entry; MgE) to collect the alcohol. Training continued until stable performance was achieved under a fixed ratio 3 (FR3) schedule, in which three active lever presses triggered an alcohol delivery. ***B***, Schematic of virus infusion for in vivo fiber photometry recordings. In D1-Cre rats, one hemisphere of the DMS was infused with AAV-FLEX-jGCaMP7f (GCaMP), and the other with AAV-GRAB_DA2m_ (DA2m). Virus assignments were counterbalanced across animals. ***C***, Sample confocal images show GCaMP expression in the cytosol of dMSNs (left) and DA2m expression on the membrane of DMS neurons (right). ***D***, Representative raw traces of calcium and dopamine signals recorded during an OSA session. ***E***, Behavioral distribution from all rats under the FR3 schedule. On average, rats entered the magazine ∼1.5 sec after the onset of alcohol delivery. ***F***, Representative heatmaps showing calcium activity during an FR3 session from one rat. The dashed lines at 0 sec and 1.5 sec indicate the timing of alcohol delivery and the average time of first magazine entry, respectively. ***G***, Averaged across all animals, dMSN calcium peaked during the transition between active pressing and magazine entry, followed by a dip at magazine entry. ***H***, Representative heatmaps showing dopamine release during an FR3 session in one rat. ***I***, Averaged across all animals, dopamine signal peaked immediately after active pressing and delivery. ***J***, The cosine similarity heatmap reveals a high correlation between the behaviors and the area-under-curve (AUC) of the dMSN GCaMP peak, as enclosed by the dashed rectangle. X-axis from left to right: *T_1st active press_*, time of the first active press; MgE half width, half width of averaged duration of magazine entry. Y-axis: P_max_, z-score of GCaMP peak; *T*(P_max_), time of GCaMP peak; P_min_, GCaMP dip; *T*(P_min_), time of GCaMP dip; AUC(P), AUC of GCaMP peak. ***K***, The AUC of the GCaMP peak (left; *t*_(11)_ = -0.01, *p* > 0.05) or the z-score of the DA peak (right; *t*_(13)_ = 1.76, *p* > 0.05) between the left and right hemisphere had no difference. Cosine similarity test (J) and unpaired t test (K). n = 11 rats (G), 5 rats (K; GCaMP-left), 8 rats (K; GCaMP-right), 7 rats (K; DA-left), 8 rats (K; DA-right).

Training began under a fixed-ratio 1 (FR1) schedule and progressed to FR3 once stable responding was established. Each alcohol delivery was followed by a 20-sec time-out period to allow consumption. Fiberphotometry was performed under the FR3 schedule. Behaviorally, animals took approximately 1.5 seconds to enter the magazine after the final active lever press (Fig. 1E). Notably, the dMSN GCaMP signal exhibited a distinct peak immediately after the last active press but before magazine entry (Fig. 1F, G; between 0 and 1.5 sec), consistent with previous reports.^38,39^ This peak quickly transitioned to a dip once the animals began to enter the magazine to collect alcohol (Fig. 1F, G; after 1.5 sec). In contrast, dopamine signals peaked in synchrony with alcohol delivery (Fig. 1H, I). Cosine similarity analysis revealed strong correlations between dMSN calcium signals (area-under-curve [AUC] of the GCaMP peak) and task-related behaviors (active pressing and magazine entry) (Fig. 1J). Therefore, we focused on the AUC of the GCaMP signal for subsequent comparisons. GCaMP and DA2m signals were comparable between hemispheres (Fig. 1K).

Together, these results show that DMS dMSNs are most active immediately following lever pressing—prior to alcohol collection—while DMS dopamine release is tightly time-locked to the delivery of alcohol.

### dMSN activity is elevated during cue-alcohol memory retrieval after abstinence but normalized by extinction training

Having established the activation pattern of dMSN and release pattern of dopamine during alcohol OSA, we next investigated whether abstinence or extinction training differently affects dMSN activity and dopamine release during cue-induced reinstatement. After ∼6 weeks of alcohol OSA training, rats were separated into a forced abstinence group, which stayed in their home-cage for 9 d, or an extinction group, which was subjected to 9 d of extinction training (Fig. 2A). This was followed by a 2-d cue-induced reinstatement test. Fiber photometry recordings were conducted during the final FR3 training sessions and reinstatement testing (Fig. 2A).

**Figure 2.**
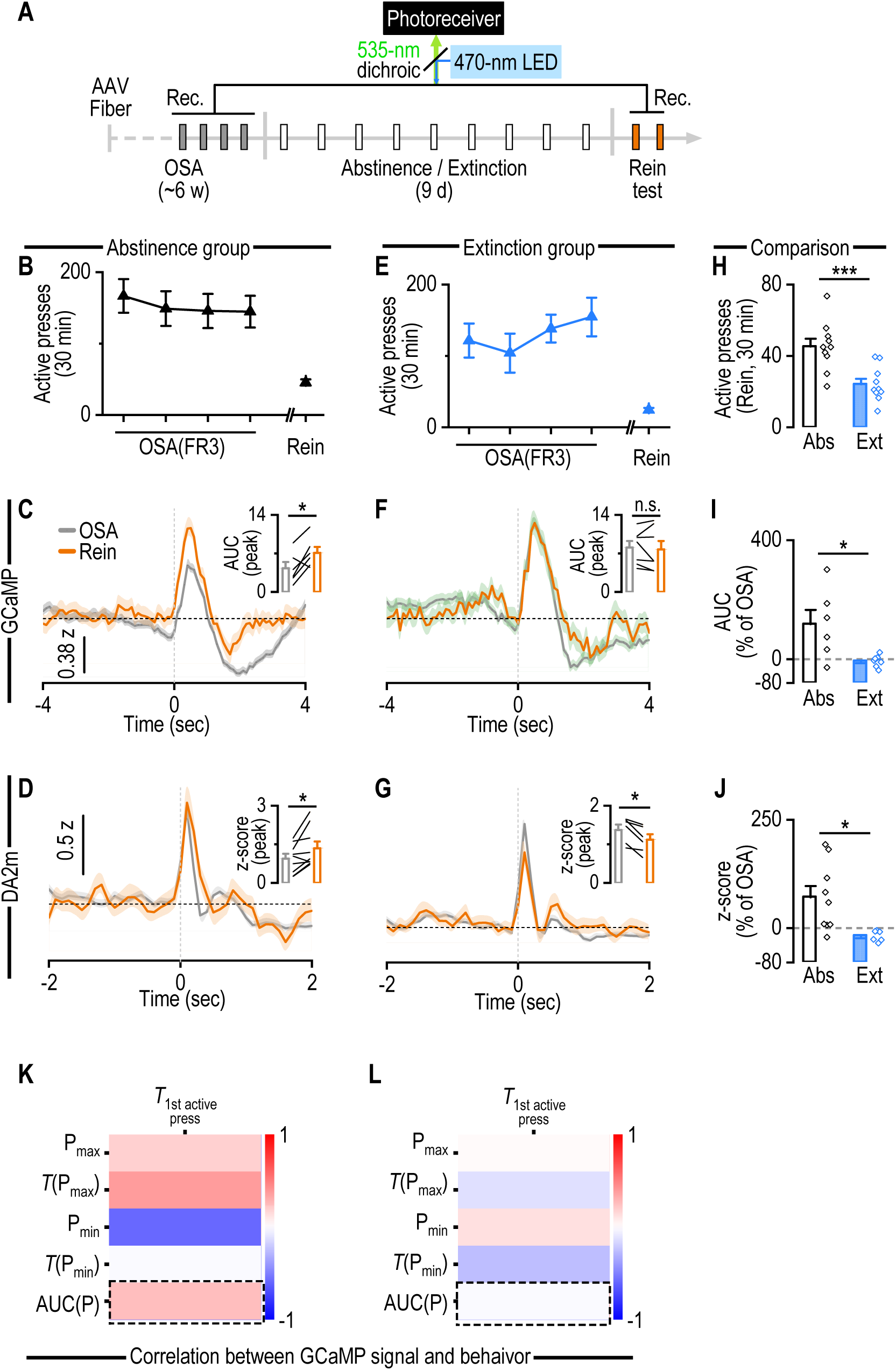
Abstinence enhances dMSN activity and dopamine release during cue-induced reinstatement, an effect that is normalized by extinction training. ***A***, Experimental timeline. D1-Cre rats underwent ∼6 weeks of alcohol OSA training under an FR3 schedule. Baseline recordings of calcium and dopamine signals were performed over the last four sessions. Rats were then assigned to either the abstinence group (Abs; 9 days of home-cage abstinence) or the extinction group (Ext; 9 days of operant lever-extinction training). Cue-induced reinstatement (Rein) tests were conducted across two days, with GCaMP and DA2m recorded on separate days. ***B***, Active lever presses during the last four OSA sessions and during reinstatement (average of two sessions) in the abstinence group. ***C***, In abstinent rats, dMSN GCaMP exhibited an increased AUC of the peak (*t*_(5)_ = -2.86, \**p* < 0.05) during reinstatement compared to the OSA. ***D***, In abstinent rats, the z-score of the DMS dopamine peak increased during reinstatement compared to the OSA. *t*_(8)_ = -2.74, \**p* < 0.05. ***E***, Active lever presses during the last four OSA sessions and reinstatement (average) in the extinction group. ***F***, No difference in GCaMP peak AUC was observed during reinstatement compared to OSA following extinction training. *t*_(6)_ = 0.519, *p* > 0.05. ***G***, After extinction, the z-score of the DMS dopamine peak reduced during reinstatement compared to the OSA. *t*_(5)_ = 3.79, \**p* < 0.05. ***H***, Extinction training significantly reduced active lever presses during reinstatement compared to abstinence. *t*_(18)_ = 3.98, \*\*\**p* < 0.001. ***I***, Percentage change in dMSN GCaMP peak AUC from OSA to reinstatement was smaller in the extinction group than in the abstinence group. *t*_(11)_ = 2.77, \**p* < 0.05. ***J***, Percentage change in dopamine peak from OSA to reinstatement was smaller in the extinction group than in the abstinence group. *t*_(13)_ = 2.71, \**p* < 0.05. ***K***,***L***, The correlation between the GCaMP signal and the first active press was still present in the abstinence group (K) but absent in the extinction group (L), as marked by dashed rectangles. Sample traces (left of C, D, F, G) were averaged data from all animals in the corresponding group during OSA or reinstatement. Paired *t* test (C, F, D, G), unpaired *t* test (H-J), and cosine similarity test (K). n = 10 rats (B), 6 rats (C), 9 rats (D), 10 rats (E), 7 rats (F), 6 rats (G), 10 rats per group (H), 6 rats (I; Abs), 7 rats (I; Ext), 9 rats (J; Abs), 6 rats (J; Ext), 6 rats (K), and 7 rats (L).

In the abstinence group (Fig. 2B), dMSN activity—measured as the GCaMP peak AUC— was significantly elevated during reinstatement compared to the FR3 baseline (Fig. 2C). Dopamine release also increased markedly during reinstatement (Fig. 2D). This suggests that dMSN activity is enhanced during the retrieval of a cue-alcohol memory after abstinence, potentially attributable to amplified dopamine release. Conversely, in the extinction group (Fig. 2E), there was no notable change in the AUC of the GCaMP peak between the FR3 baseline and reinstatement (Fig. 2F). Intriguingly, dopamine signals exhibited a marked decrease during reinstatement (Fig. 2G). These results indicated that extinction training dampened dopamine release, which could consequently alter corticostriatal plasticity and normalize dMSN activity during memory retrieval.

Further comparison between the abstinence and extinction groups revealed that the extinction group had significantly fewer active presses during reinstatement (Fig. 2H). The increase in dMSN activity from baseline to reinstatement was significantly lower in the extinction group (Fig. 2I), as was the change in dopamine release (Fig. 2J). Notably, GCaMP signals remained correlated with lever pressing in the abstinence group (Fig. 2K), but this association was weaker following extinction training (Fig. 2L).

Overall, these findings suggest that abstinence enhances dMSN activity and dopamine release during cue-induced relapse, while extinction training reduces both, potentially mitigating relapse vulnerability.

### Optogenetic induction of mPFC-to-dMSN synaptic plasticity after abstinence or extinction oppositely controls cue-alcohol memory retrieval

Having demonstrated that abstinence and extinction training differentially modulate dMSN activity and dopamine release during cue-induced reinstatement, we next sought to elucidate the underlying mechanism connecting these physiological changes to relapse behaviors. Because dopamine release critically modulates corticostriatal synaptic plasticity onto dMSNs,^21,28,32,40^ the observed alterations in dopamine signaling and dMSN activity may reflect underlying changes in corticostriatal synaptic strength. Therefore, we hypothesized that directly manipulating corticostriatal plasticity—specifically, inducing LTD after abstinence to mimic extinction-like plasticity, or inducing LTP after extinction to mimic abstinence-like potentiation—would bidirectionally regulate the propensity for cue-induced relapse.

To test this hypothesis, we aimed to experimentally alter corticostriatal plasticity following periods of abstinence or extinction training and assess the subsequent impact on cue-induced reinstatement. D1-Cre rats received bilateral infusions of AAV-Chronos-GFP in the mPFC and AAV-FLEX-Chrimson-tdTomato in the DMS, with optical fibers implanted bilaterally above the DMS (Fig. 3A, 3B). Rats were trained to self-administer alcohol in operant chambers until stable performance was achieved under an FR3 schedule. Subsequently, the rats were categorized into abstinence or extinction groups and subjected to a cue-induced reinstatement test (Fig. 3C).

**Figure 3.**
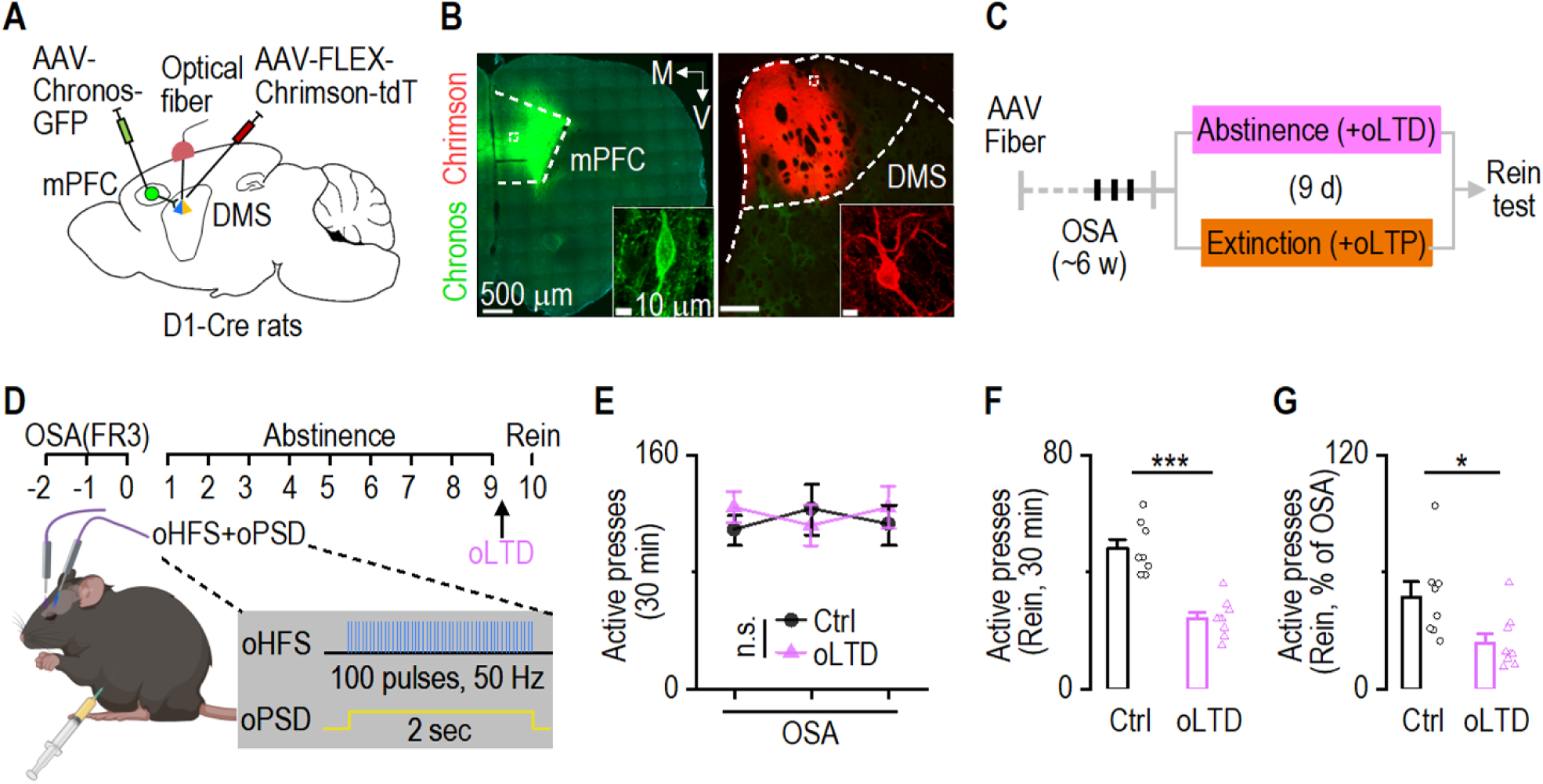
mPFC-to-dMSN LTD after abstinence reduces cue-induced reinstatement. ***A***, Schematic illustrating virus infusion and fiber implantation in D1-Cre rats. ***B***, Representative confocal images of AAV-Chronos-GFP expression in the mPFC (left) and AAV-FLEX-Chrimson-tdTomato expression in the DMS (right). The insets show single neurons from enclosed areas. M, medial; V, ventral. ***C***, Experiment timeline. After ∼6 weeks of alcohol OSA training, rats were divided into abstinence or extinction groups. On the last day of abstinence or extinction, we delivered optogenetic long-term depression (oLTD) or long-term potentiation (oLTP) induction protocols, followed by a cue-induced reinstatement test the subsequent day. ***D***, The schematic illustrating the schedule and protocol for oLTD induction in the abstinence group. ***E-G***, The oLTD and control (Ctrl) groups exhibited similar levels of active presses during the OSA (E; *F*_(1,15)_ = 0.03, *p* > 0.05). However, oLTD induction decreased the number of active presses (F; *t*_(15)_ = 6.46, \*\*\**p* < 0.001) and normalized active presses (G, normalized to the OSA; *t*_(15)_ = 2.64, \**p* < 0.05) during the reinstatement. Two-way RM ANOVA followed by Tukey *post hoc* test (E), unpaired *t* test (F, G). n = 8 rats (Ctrl) and 9 rats (oLTD).

In the abstinence group, animals remained in their home-cages for 9 d. On the final day of abstinence, corticostriatal LTD was induced in a neutral chamber via pre-injection of the NMDA receptor antagonist MK801 (i.p., 0.1 mg/kg) 15 min before the pairing of oHFS and oPSD,^32^ aiming to depotentiate corticostriatal plasticity and mimic the effect of extinction (Fig. 3D). Control animals in the abstinence group did not receive LTD induction. Previous studies confirmed that MK801 administration itself did not alter corticostriatal plasticity nor alcohol-seeking behaviors in rats.^32^ Although both the control and optogenetically-induced LTD (oLTD) groups displayed similar active presses during FR3 (Fig. 3E), rats in the oLTD group exhibited significantly fewer active presses during the reinstatement test (Fig. 3F). This difference persisted even when the active presses were normalized to those during the baseline FR3 sessions (Fig. 3G). These findings demonstrate that depotentiation of corticostriatal plasticity following abstinence effectively mimics the behavioral outcomes of extinction training, reducing relapse vulnerability.

In the extinction group, animals underwent extinction training in the operant chamber for 9 d. After the last extinction training, corticostriatal LTP was induced in a neutral chamber by pairing oHFS with oPSD (Fig. 4A), with the aim of potentiating corticostriatal plasticity and mimic abstinence. Rats in both the control and optogenetically-induced LTP (oLTP) groups displayed similar performance during baseline FR3 and extinction training (Fig. 4B). However, rats subjected to oLTP showed a pronounced increase in active presses during the reinstatement test (Fig. 4C). This effect persisted even after normalizing the presses to those seen during the baseline FR3 sessions (Fig. 4D). These results indicate that experimentally potentiating corticostriatal synapses after extinction training partially reverses the extinction effect and facilitates relapse.

**Figure 4.**
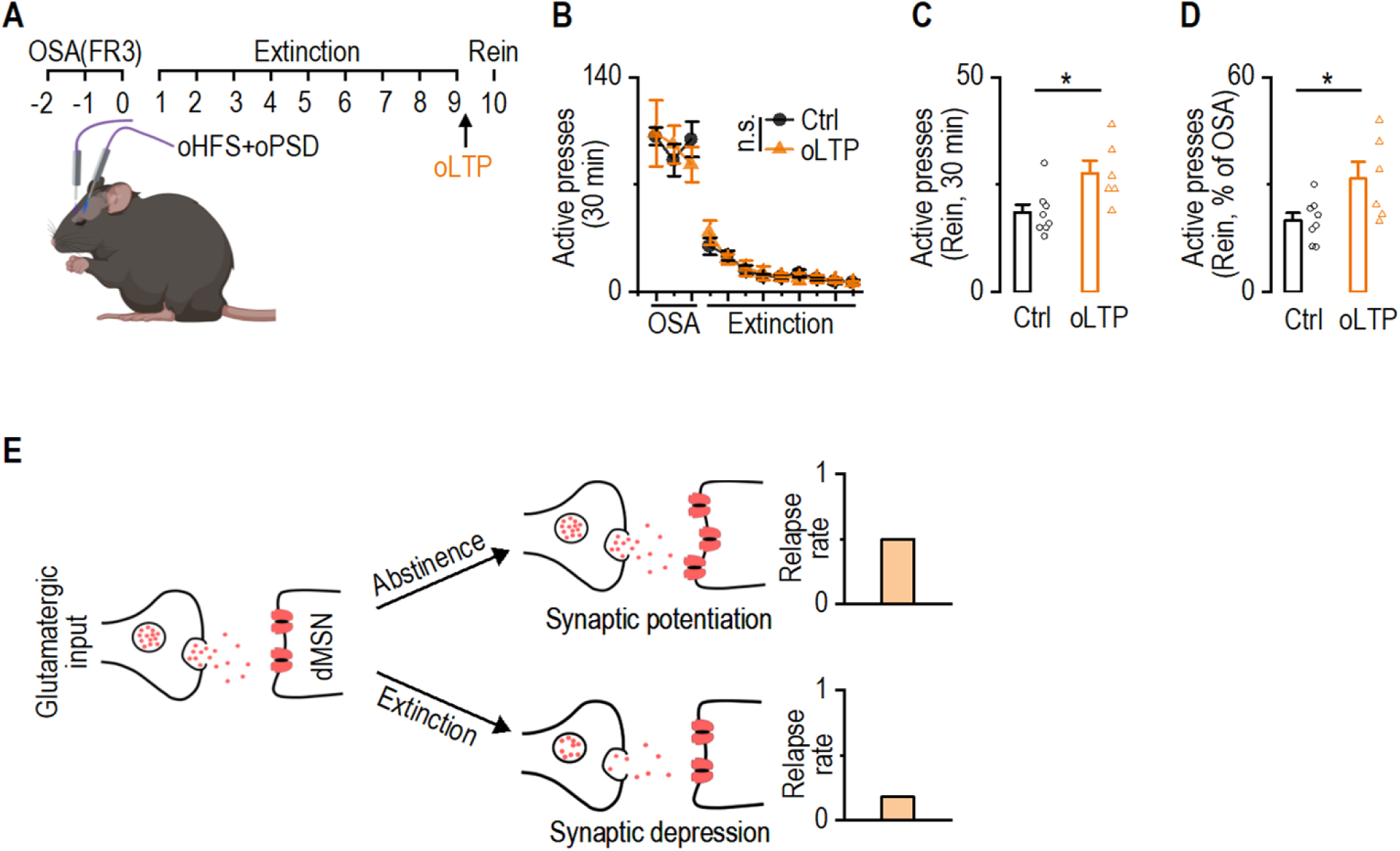
mPFC-to-dMSN LTP after extinction potentiates cue-induced reinstatement. ***A***, The schematic outlining the schedule and protocol for oLTP induction following 9 d of extinction training. ***B***-***D***, The oLTP and Ctrl groups exhibited similar levels of active presses during the OSA and extinction training (B; *F*_(1,12)_ = 0.05, *p* > 0.05). However, oLTP induction increased the number of active presses (C; *t*_(12)_ = -2.74, \**p* < 0.05) and normalized active presses (D, normalized to the OSA; *t*_(15)_ = -2.46, \**p* < 0.05) during reinstatement. ***E***, Schematic of the working hypothesis: glutamatergic transmission onto DMS dMSNs is strengthened during abstinence, promoting relapse; conversely, extinction weakens this transmission, reducing relapse. Two-way RM ANOVA followed by Tukey *post hoc* test (B), unpaired *t* test (C, D). n = 8 rats (Ctrl) and 6 rats (oLTP).

In summary, our fiber photometry results suggest that abstinence enhances dopamine release and dMSN activity during cue-induced relapse, whereas extinction training reduces dopamine signaling and normalizes dMSN activity. Although we did not directly measure synaptic changes, these physiological patterns are consistent with a model in which corticostriatal plasticity is differentially shaped by abstinence and extinction history. Specifically, the increase in dopamine during abstinence may potentiate mPFC-to-dMSN synapses, thereby facilitating relapse behavior (Fig. 4E). In contrast, reduced dopamine signaling following extinction may weaken these synapses, contributing to relapse suppression (Fig. 4E). Our optogenetic manipulations support this interpretation by demonstrating that mimicking synaptic potentiation or depression can bidirectionally influence relapse vulnerability.

## Discussion

In this study, we identified distinct neural signatures and synaptic mechanisms underlying alcohol relapse after abstinence or extinction, and demonstrated that bidirectional modulation of corticostriatal synaptic strength causally controls cue-induced reinstatement of alcohol-seeking behavior. Using in vivo calcium and dopamine recordings, we found that abstinence enhances DMS dMSN activity and dopamine release during relapse, whereas extinction training normalizes dMSN activity and reduces dopamine release. Importantly, manipulating mPFC-to-dMSN synaptic plasticity was sufficient to either attenuate or reinstate relapse-like behavior, revealing a causal role for corticostriatal dynamics in the expression of alcohol-associated memories.

Our fiber photometry recordings, conducted in freely-moving rats during an operant task, provide compelling in vivo evidence of neural activity linked to behavior. The task used reflects a classic operant conditioning paradigm,^37,41^ in which active lever pressing (reward-seeking) leads to alcohol delivery, followed by magazine entry (reward-taking). dMSNs are known to be involved in action initiation,^42^ but in our paradigm, we observed that dMSN activity did not precede the start of an action sequence (i.e., active lever press). Instead, activity dipped during the lever presses and peaked at the transition from pressing to magazine entry, suggesting a role in goal evaluation or outcome expectation rather than movement initiation.^43–45^ This aligns with prior studies showing DMS activation during reward delivery rather than action onset.^39^ Notably, similar activation patterns have been observed in food-reward paradigms, where peak striatal activity corresponds with the movement toward reward collection, and not with movements of similar motor demands like lever pressing.^38^ Importantly, this population-level activation of dMSNs may also suggest the formation of neuronal ensembles that encode the learned association between lever-press actions and alcohol reward.^46^ Dopamine release exhibited a concurrent or slightly earlier peak, ramping up during lever pressing and peaking after reward delivery. This signal likely reflects reward expectation and valuation,^47–49^ potentially integrating multisensory cues^48^ (e.g., sound of dipper, cue light, smell of alcohol) and reinforcing synaptic connections between active glutamatergic inputs (e.g., mPFC) and dMSNs. Given that dopamine is a key modulator of corticostriatal plasticity,^21,28,29,48,50^ and that memory is thought to be encoded by synaptic changes,^51^ this convergence of dopamine release and dMSN activity may facilitate the formation and stabilization of alcohol-associated corticostriatal plasticity that encodes operant associative memories, ultimately driving later relapse.

Our result that increased activity in dMSNs during cue-induced reinstatement of alcohol seeking following a 9-day abstinence reinforces previous findings of heightened striatal neuron activity during drug relapse.^52–54^ Cue-induced dopamine release in the dorsal striatum is a well-documented phenomenon across multiple drugs,^66,67^ including alcohol. Recent work even shows that re-exposure to cocaine cues after abstinence amplifies dopamine signaling in the pallidum.^55^ In our study, similar cue-induced dopamine elevations were observed in the DMS following abstinence, potentially driving increased dMSN activity. Additionally, alcohol-induced glutamate spillover—due to impaired perisynaptic astrocytic transporter function—could prolong cue-evoked glutamatergic signaling,^56^ further enhancing dMSN activation and potentially recruiting new ensemble members. These mechanisms may underlie the incubation of craving, a phenomenon wherein cue-induced drug seeking intensifies over time following abstinence.^6^

Conversely, extinction training appears to dampen the neural processes supporting relapse. Extinction is known to share features with synaptic depression and to impair plasticity induction.^15,57,58^ Previous studies have shown that extinction can occlude LTD in the amygdala and prevent LTD induction in ventrostriatal dMSNs.^15,57,59^ These findings support the idea that extinction weakens memory retrieval by disrupting corticostriatal synaptic efficacy. We found that diminished dopamine release during cue-induced relapse after extinction may reflect reduced reward prediction and contribute to synaptic weakening, ultimately decreasing dMSN recruitment and relapse behavior.

Critically, we provide causal support for this framework through optogenetic manipulation of corticostriatal plasticity. Inducing LTD after abstinence reduced relapse, likely by depotentiating alcohol-strengthened synapses.^32,33^ This effect may be mediated through presynaptic mechanisms that lower glutamate release,^32^ thereby decreasing dMSN activation. Conversely, inducing LTP after extinction reversed the protective effect of extinction, promoting relapse through potentiation of mPFC-to-dMSN synaptic transmission. These results demonstrate that corticostriatal plasticity is not only shaped by behavioral history but can also be targeted to manipulate relapse vulnerability.

Together, our findings reveal a mechanistic framework in which abstinence and extinction differentially shape striatal circuit dynamics and dopamine signaling, and where mPFC-to-dMSN synaptic plasticity acts as a bidirectional switch controlling alcohol relapse. This work advances our understanding of relapse neurobiology and identifies specific synaptic processes that may be targeted to enhance the durability of extinction and reduce the risk of relapse in alcohol addiction.

## Methods

### Reagents

AAV9-Syn-FLEX-jGCaMP7f (#104491) and AAV9-hSyn-GRAB_DA2m (#140553) were obtained from Addgene. AAV8-Syn-Chronos-GFP and AAV8-Syn-FLEX-Chrimson-tdTomato were purchased from the University of North Carolina Vector Core. MK801 were purchased from Sigma.

### Animals

D1-Cre rats were obtained from Rat Resource and Research Center.^60^ Genotypes were determined by PCR analysis of Cre gene in tail DNA.^18,61,62^ All animals that were used are in mixed gender and aged from 3-8 months. Animals were housed in a temperature- and humidity-controlled vivarium with a reversed 12-h light/dark cycle (lights on at 10:00 P.M. and off at 10:00 A.M.). All behavior experiments were conducted in their dark cycle, starting approximately 1 h after the light went off. Food and water were available *ad libitum*. All animal care and experimental procedures were approved by the Texas A&M University Institutional Animal Care and Use Committee.

### Experiment procedures

#### Intermittent-access to 20% alcohol two-bottle-choice drinking procedure

To acclimate rats with alcohol and establish high levels of alcohol consumption, we utilized the intermittent-access to 20% alcohol two-bottle-choice drinking procedure.^18,23,31,33,63^ D1-Cre rats were given free access to two bottles containing water or 20% alcohol for three 24-h sessions (Mondays, Wednesdays, and Fridays), with 24-h or 48-h withdrawal periods (Tuesdays, Thursdays, Saturdays, and Sundays) each week. During the withdrawal periods, the rats had unlimited access to water bottles. This procedure was followed for ∼3 weeks.

### Operant self-administration of alcohol

Animals were trained to self-administer 20% alcohol in operant chambers using a fixed ratio (FR) schedule, as described previously.^30,32,64^ The rat operant system (Coulbourn Instrument) include two levers: active and inactive in each chamber, on the opposite sides of the same wall.

During training sessions, a house light centered above the operant illuminated. Each operant experiment commenced with continuous reinforcement (FR1) for approximately two to three weeks, wherein pressing the active side resulted in a 0.1 mL delivery of 20% alcohol. Actions on the inactive side were documented but did not initiate a programmed event. The alcohol solution was dispensed into a stainless steel reservoir situated within the magazine port between active and inactive levers. Each alcohol delivery persisted for 20 sec, during which the magazine port was lit by a discrete yellow cue light for the same duration; both levers were withdrawn during the delivery period to facilitate alcohol consumption. When animals were able to achieve at least 10 deliveries under the FR1 schedule, the criterion to receive the alcohol escalated to FR2 (around one to two weeks), eventually progressing to FR3 (around two weeks). Each operant session for rats lasted 30 min.

#### Extinction training

The extinction training was conducted under the FR3 schedule. However, pressing on the active side did not result in actual alcohol delivery, nor cue light presentation. Each extinction session lasted 30 min, and in total for 9 d.

#### Cue-induced reinstatement of alcohol seeking

The reinstatement test was conducted as described.^65^ It was carried out at the FR3 schedule, where three actions on the active side triggered the illumination of the cue light within the magazine port and the dipper action sound, with the exception that the first cue presentation and dipper action was immediately after the first active press. Considering that many rats stopped active responses following the 9-day extinction training, a 20 µL volume of alcohol was pipetted into the magazine port prior to the session’s commencement to instigate the animals’ responses. However, once the session had started, alcohol was not available.

### Stereotaxic virus infusion and fiber implantation

The stereotaxic virus infusion procedure was conducted as described previously.^32,33,62,66^ Where required for the experimental design, AAV-Chronos-GFP (0.8 µL/site) was infused into the mPFC (AP: +2.9 mm, ML: ±0.65 mm, DV: -3.5 mm from the Bregma), AAV-FLEX-Chrimson-tdT (1.2 µL/site) was infused into the DMS (AP: +0.36 mm, ML: ±2.3 mm, DV: -4.7 mm from the Bregma). For fiberphotometry rats, one side of DMS was infused with AAV-FLEX-jGCaMP7f (1.2 µL) and the other side with AAV-GRAB_DA2m (1.2 µL)(AP: +0.36 mm, ML: ±2.3 mm, DV: -4.7 mm from the Bregma; the assignment of jGCaMP7f and GRAB_DA2m was counterbalanced). Optical fiber implants (300-μm diameter optical fiber secured to a 2.5-mm stainless steel ferrule) were placed bilaterally in DMS through the injection tract (DV: -4.6 mm). Animals were anesthetized with 3-4% isoflurane at 1.0 L/min and mounted in a stereotaxic surgery frame. The head was leveled and craniotomy was performed using stereotaxic coordinates adapted from the brain atlas.^67^ The virus was infused at a rate of 0.08 µL/min. At the end of the infusion, the injectors remained at the injection site for an additional 10-15 min before removal to allow for virus diffusion. For fiber implants, four metal screws were fixed into the skull to support the implants, which were further secured with dental cement (Henry Schein). The scalp incision was then sutured, and the animals were returned to their home cage for recovery.

### In vivo fiber-photometry recording

Fiberphotometry was conducted as previously described.^38,68^ In each D1-Cre rats, one side of the DMS contained FLEX-jGCaMP7f and the other side contained GRAB_DA2m. The left or right assignment of these two sensors were counterbalanced. Rats were connected with fiber for the entire training periods to acclimate to the fiber patch cable. Baseline recordings occurred during the last 4 sessions of FR3 training (each sensor in each animal was recorded twice and then the signals were averaged). Following 9-d extinction training or abstinence, cue-induced reinstatement was tested for 2 sessions (1 session for GCaMP measurement and 1 session for DA2m measurement; the measurement was counterbalanced). The spectrum data was recorded continuously at 10 Hz. At the same time, behavior data were collected. Fiber-photometry data was collected using OceanView 1.6.7. 488 nm laser was delivered to excite jGCaMP7f expressing in dMSNs or GRAB_DA2m. The Z-score (Z) was calculated by

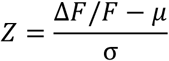

using MATLAB. To analyze the fiberphotometry data in the context of rat behavior, MATLAB scripts were developed. Cosine similarity between the behavior data and the photometry data was analyzed, and the Pearson correlation coefficients (\rho) between the behavior data (x) and the fiber photometry data (y) is calculated by

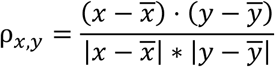

using MATLAB. Both the extinction and abstinence groups had 10 rats. However, due to recording errors or fiber misplacement, 4 recordings of GRAB_DA2m and 3 recordings of jGCaMP7f were missed in the extinction group; 1 recording of GRAB_DA2m and 4 recordings of jGCaMP7f were missed in the abstinence group.

### In vivo LTP and LTD induction

During the last day of abstinence or after the last session of extinction training, D1-Cre rats received an LTD/LTP-inducing protocol in a neutral Plexiglass chamber, with no visual cues. LTP induction consists of paired oHFS + oPSD using the following protocol: 100 pulses at 50 Hz of 473-nm light (2 ms) with constant 590-nm light for 2 sec. Each stimulation was repeated 7 times with a 20-sec interval, forming a bout of stimulation. A total of 6 bouts were delivered with a 3-min interval. LTD induction employed the following protocol: animals were injected with MK801 (0.1 mg/kg) 15 min before delivery of oHFS and oPSD (same as above). Control animals for both inductions were exposed to the Plexiglass chamber and connected to the patch cable, but no light was delivered. The complete LTP/LTD-inducing procedure was performed once, and animals were tested for cue-induced reinstatement 24 h later.

### Histology

Animals were perfused intracardially with 4% formaldehyde in phosphate-buffered saline. The brains were removed and post-fixed overnight in 4% formaldehyde in phosphate-buffered saline at 4°C prior to dehydration in 30% sucrose solution. The brains were cut into 50-µm coronal sections using a cryostat. All images were acquired using a confocal microscope (FluoView 3000, Olympus, Tokyo, Japan) and analyzed using IMARIS 8.3.1 (Bitplane, Zürich, Switzerland), as previously reported.^61,62^

### Statistical analysis

All data are expressed as the mean ± the standard error of the mean. Data were analyzed by two-tailed *t* test (unpaired or paired) and two-way ANOVA with repeated measurement, followed by the Tukey post hoc test. Significance was determined if *p* < 0.05. Statistical analysis was conducted by the SigmaPlot program. Graphs were constructed using the OriginPro program.

## Acknowledgements

This research was supported by NIAAA Grant R01AA021505, R01AA027768, U01AA025932 and X-grant from Texas A&M University to J.W., McGovern Fellowship from Texas Research Society on Alcoholism (TRSA) and Doctoral Student Small Grant from Research Society on Alcohol (RSA) to X.X.

## Declaration of interests

The authors declare no competing interests.

## Inclusion and diversity

We support inclusive, diverse, and equitable conduct of research.

## Author contributions

Conceptualization: X.X., J.W.; Methodology: X.X., R.C., J.W.; Investigation and Formal analysis: X.X., R.C.; Software: R.C.; Writing - Original Draft: X.X.; Writing – Review & Editing: J.W., X.X; Resources: X.W.; Funding acquisition: J.W., and X.X.; Supervision: J.W.

## Notes

### Competing Interest Statement

The authors have declared no competing interest.

## References

1. Torregrossa, M.M., Corlett, P.R., and Taylor, J.R. (2011). Aberrant learning and memory in addiction. Neurobiology of learning and memory 96, 609–623. 10.1016/j.nlm.2011.02.014.

2. Nicholas W. Gilpin, P.D., and George F. Koob, Ph.D. (2008). Neurobiology of Alcohol Dependence. Alcohol research & health : the journal of the National Institute on Alcohol Abuse and Alcoholism.

3. Connor, J.P., Haber, P.S., and Hall, W.D. (2016). Alcohol use disorders. Lancet 387, 988–998. 10.1016/S0140-6736(15)00122-1.

4. Heilig, M., Egli, M., Crabbe, J.C., and Becker, H.C. (2010). Acute withdrawal, protracted abstinence and negative affect in alcoholism: are they linked? Addiction biology 15, 169–184. 10.1111/j.1369-1600.2009.00194.x.

5. Bienkowski, P., Rogowski, A., Korkosz, A., Mierzejewski, P., Radwanska, K., Kaczmarek, L., Bogucka-Bonikowska, A., and Kostowski, W. (2004). Time-dependent changes in alcohol-seeking behaviour during abstinence. Eur Neuropsychopharmacol 14, 355–360. 10.1016/j.euroneuro.2003.10.005.

6. Pickens, C.L., Airavaara, M., Theberge, F., Fanous, S., Hope, B.T., and Shaham, Y. (2011). Neurobiology of the incubation of drug craving. Trends Neurosci. 34, 411–420. 10.1016/j.tins.2011.06.001.

7. Drummond, D.C., Cooper, T., and Glautier, S.P. (1990). Conditioned learning in alcohol dependence: implications for cue exposure treatment. Br J Addict 85, 725–743. 10.1111/j.1360-0443.1990.tb01685.x.

8. Drummond, D.C., and Glautier, S. (1994). A controlled trial of cue exposure treatment in alcohol dependence. Journal of consulting and clinical psychology 62, 809–817. 10.1037//0022-006x.62.4.809.

9. Torregrossa, M.M., and Taylor, J.R. (2013). Learning to forget: manipulating extinction and reconsolidation processes to treat addiction. Psychopharmacology (Berl) 226, 659–672. 10.1007/s00213-012-2750-9.

10. Vollstadt-Klein, S., Loeber, S., Kirsch, M., Bach, P., Richter, A., Buhler, M., von der Goltz, C., Hermann, D., Mann, K., and Kiefer, F. (2011). Effects of cue-exposure treatment on neural cue reactivity in alcohol dependence: a randomized trial. Biological psychiatry 69, 1060–1066. 10.1016/j.biopsych.2010.12.016.

11. Milton, A.L., and Everitt, B.J. (2012). The persistence of maladaptive memory: addiction, drug memories and anti-relapse treatments. Neurosci Biobehav Rev 36, 1119–1139. 10.1016/j.neubiorev.2012.01.002.

12. Cofresi, R.U., Lewis, S.M., Chaudhri, N., Lee, H.J., Monfils, M.H., and Gonzales, R.A. (2017). Postretrieval Extinction Attenuates Alcohol Cue Reactivity in Rats. Alcohol Clin Exp Res 41, 608–617. 10.1111/acer.13323.

13. Ray, L.A., Grodin, E.N., Leggio, L., Bechtholt, A.J., Becker, H., Feldstein Ewing, S.W., Jentsch, J.D., King, A.C., Mason, B.J., O’Malley, S., et al. (2021). The future of translational research on alcohol use disorder. Addiction biology 26, e12903. 10.1111/adb.12903.

14. Roberts-Wolfe, D., Bobadilla, A.C., Heinsbroek, J.A., Neuhofer, D., and Kalivas, P.W. (2018). Drug Refraining and Seeking Potentiate Synapses on Distinct Populations of Accumbens Medium Spiny Neurons. J. Neurosci. 38, 7100–7107. 10.1523/Jneurosci.0791-18.2018.

15. Giannotti, G., Gong, S., Fayette, N., Heinsbroek, J.A., Orfila, J.E., Herson, P.S., Ford, C.P., and Peters, J. (2021). Extinction blunts paraventricular thalamic contributions to heroin relapse. Cell reports 36, 109605. 10.1016/j.celrep.2021.109605.

16. Yin, H.H., Ostlund, S.B., Knowlton, B.J., and Balleine, B.W. (2005). The role of the dorsomedial striatum in instrumental conditioning. Eur J Neurosci 22, 513–523. 10.1111/j.1460-9568.2005.04218.x.

17. Balleine, B.W., Delgado, M.R., and Hikosaka, O. (2007). The role of the dorsal striatum in reward and decision-making. J Neurosci 27, 8161–8165. 10.1523/JNEUROSCI.1554-07.2007.

18. Wang, J., Cheng, Y., Wang, X., Roltsch Hellard, E., Ma, T., Gil, H., Ben Hamida, S., and Ron, D. (2015). Alcohol Elicits Functional and Structural Plasticity Selectively in Dopamine D1 Receptor-Expressing Neurons of the Dorsomedial Striatum. J Neurosci 35, 11634–11643. 10.1523/JNEUROSCI.0003-15.2015.

19. Wang, J., Ben Hamida, S., Darcq, E., Zhu, W.H., Gibb, S.L., Lanfranco, M.F., Carnicella, S., and Ron, D. (2012). Ethanol-Mediated Facilitation of AMPA Receptor Function in the Dorsomedial Striatum: Implications for Alcohol Drinking Behavior. J. Neurosci. 32, 15124–15132. 10.1523/Jneurosci.2783-12.2012.

20. Lu, J., Cheng, Y., Xie, X., Woodson, K., Bonifacio, J., Disney, E., Barbee, B., Wang, X., Zaidi, M., and Wang, J. (2021). Whole-Brain Mapping of Direct Inputs to Dopamine D1 and D2 Receptor-Expressing Medium Spiny Neurons in the Posterior Dorsomedial Striatum. eNeuro 8, ENEURO.0348-0320.2020. 10.1523/ENEURO.0348-20.2020.

21. Gerfen, C.R., and Surmeier, D.J. (2011). Modulation of striatal projection systems by dopamine. Annu Rev Neurosci 34, 441–466. 10.1146/annurev-neuro-061010-113641.

22. Kravitz, A.V., Tye, L.D., and Kreitzer, A.C. (2012). Distinct roles for direct and indirect pathway striatal neurons in reinforcement. Nat Neurosci 15, 816–818. 10.1038/nn.3100.

23. Cheng, Y., Huang, C.C.Y., Ma, T., Wei, X., Wang, X., Lu, J., and Wang, J. (2017). Distinct Synaptic Strengthening of the Striatal Direct and Indirect Pathways Drives Alcohol Consumption. Biological psychiatry 81, 918–929. 10.1016/j.biopsych.2016.05.016.

24. Hong, S.I., Kang, S., Chen, J.F., and Choi, D.S. (2019). Indirect Medium Spiny Neurons in the Dorsomedial Striatum Regulate Ethanol-Containing Conditioned Reward Seeking. J Neurosci 39, 7206–7217. 10.1523/JNEUROSCI.0876-19.2019.

25. Isett, B.R., Nguyen, K.P., Schwenk, J.C., Yurek, J.R., Snyder, C.N., Vounatsos, M.V., Adegbesan, K.A., Ziausyte, U., and Gittis, A.H. (2023). The indirect pathway of the basal ganglia promotes transient punishment but not motor suppression. Neuron. 10.1016/j.neuron.2023.04.017.

26. Luscher, C., and Ungless, M.A. (2006). The mechanistic classification of addictive drugs. PLoS Med 3, e437. 10.1371/journal.pmed.0030437.

27. Maltese, M., March, J.R., Bashaw, A.G., and Tritsch, N.X. (2021). Dopamine differentially modulates the size of projection neuron ensembles in the intact and dopamine-depleted striatum. Elife 10, e68041. 10.7554/eLife.68041.

28. Bromberg-Martin, E.S., Matsumoto, M., and Hikosaka, O. (2010). Dopamine in motivational control: rewarding, aversive, and alerting. Neuron 68, 815–834. 10.1016/j.neuron.2010.11.022.

29. Surmeier, D.J., Shen, W., Day, M., Gertler, T., Chan, S., Tian, X., and Plotkin, J.L. (2010). The role of dopamine in modulating the structure and function of striatal circuits. Progress in brain research 183, 149–167. 10.1016/S0079-6123(10)83008-0.

30. Roltsch Hellard, E., Binette, A., Zhuang, X., Lu, J., Ma, T., Jones, B., Williams, E., Jayavelu, S., and Wang, J. (2019). Optogenetic control of alcohol-seeking behavior via the dorsomedial striatal circuit. Neuropharmacology 155, 89–97. 10.1016/j.neuropharm.2019.05.022.

31. Ma, T., Barbee, B., Wang, X., and Wang, J. (2017). Alcohol induces input-specific aberrant synaptic plasticity in the rat dorsomedial striatum. Neuropharmacology 123, 46–54. 10.1016/j.neuropharm.2017.05.014.

32. Ma, T., Cheng, Y., Roltsch Hellard, E., Wang, X., Lu, J., Gao, X., Huang, C.C.Y., Wei, X.Y., Ji, J.Y., and Wang, J. (2018). Bidirectional and long-lasting control of alcohol-seeking behavior by corticostriatal LTP and LTD. Nat Neurosci 21, 373–383. 10.1038/s41593-018-0081-9.

33. Xie, X., Lu, J., Ma, T., Cheng, Y., Woodson, K., Bonifacio, J., Bego, K., Wang, X., and Wang, J. (2023). Linking input- and cell-type-specific synaptic plasticity to the reinforcement of alcohol-seeking behavior. Neuropharmacology 237, 109619. 10.1016/j.neuropharm.2023.109619.

34. Bobadilla, A.C., Dereschewitz, E., Vaccaro, L., Heinsbroek, J.A., Scofield, M.D., and Kalivas, P.W. (2020). Cocaine and sucrose rewards recruit different seeking ensembles in the nucleus accumbens core. Molecular psychiatry 25, 3150–3163. 10.1038/s41380-020-00888-z.

35. Wang, W., Xie, X., Zhuang, X., Huang, Y., Tan, T., Gangal, H., Huang, Z., Purvines, W., Wang, X., Stefanov, A., et al. (2023). Striatal mu-opioid receptor activation triggers direct-pathway GABAergic plasticity and induces negative affect. Cell reports, 112089. 10.1016/j.celrep.2023.112089.

36. Panlilio, L.V., and Goldberg, S.R. (2007). Self-administration of drugs in animals and humans as a model and an investigative tool. Addiction (Abingdon, England) 102, 1863–1870. 10.1111/j.1360-0443.2007.02011.x.

37. Gardner, E.L. (2000). What we have learned about addiction from animal models of drug self-administration. Am J Addiction 9, 285–313. Doi 10.1080/105504900750047355.

38. Cui, G., Jun, S.B., Jin, X., Pham, M.D., Vogel, S.S., Lovinger, D.M., and Costa, R.M. (2013). Concurrent activation of striatal direct and indirect pathways during action initiation. Nature 494, 238–242. 10.1038/nature11846.

39. Fanelli, R.R., Klein, J.T., Reese, R.M., and Robinson, D.L. (2013). Dorsomedial and dorsolateral striatum exhibit distinct phasic neuronal activity during alcohol self-administration in rats. Eur J Neurosci 38, 2637–2648. 10.1111/ejn.12271.

40. Allichon, M.C., Ortiz, V., Pousinha, P., Andrianarivelo, A., Petitbon, A., Heck, N., Trifilieff, P., Barik, J., and Vanhoutte, P. (2021). Cell-Type-Specific Adaptions in Striatal Medium-Sized Spiny Neurons and Their Roles in Behavioral Responses to Drugs of Abuse. Frontiers in synaptic neuroscience 13, 799274. 10.3389/fnsyn.2021.799274.

41. Sanchis-Segura, C., and Spanagel, R. (2006). Behavioural assessment of drug reinforcement and addictive features in rodents: an overview. Addiction biology 11, 2–38. 10.1111/j.1369-1600.2006.00012.x.

42. Tecuapetla, F., Jin, X., Lima, S.Q., and Costa, R.M. (2016). Complementary Contributions of Striatal Projection Pathways to Action Initiation and Execution. Cell 166, 703–715. 10.1016/j.cell.2016.06.032.

43. Lau, B., and Glimcher, P.W. (2008). Value representations in the primate striatum during matching behavior. Neuron 58, 451–463. 10.1016/j.neuron.2008.02.021.

44. Shin, J.H., Kim, D., and Jung, M.W. (2018). Differential coding of reward and movement information in the dorsomedial striatal direct and indirect pathways. Nature communications 9, 404. 10.1038/s41467-017-02817-1.

45. Kwak, S., and Jung, M.W. (2019). Distinct roles of striatal direct and indirect pathways in value-based decision making. Elife 8. 10.7554/eLife.46050.

46. Cruz, F.C., Rubio, F.J., and Hope, B.T. (2015). Using c-fos to study neuronal ensembles in corticostriatal circuitry of addiction. Brain research 1628, 157–173. 10.1016/j.brainres.2014.11.005.

47. Watabe-Uchida, M., Eshel, N., and Uchida, N. (2017). Neural Circuitry of Reward Prediction Error. Annu Rev Neurosci 40, 373–394. 10.1146/annurev-neuro-072116-031109.

48. Cox, J., and Witten, I.B. (2019). Striatal circuits for reward learning and decision-making. Nat Rev Neurosci 20, 482–494. 10.1038/s41583-019-0189-2.

49. van Elzelingen W, G.J., Warnaar P, Denys D, Arbab T, Willuhn I (2022). A unidirectional but not uniform striatal landscape of dopamine signaling for motivational stimuli. Proc Natl Acad Sci U S A 119, e2117270119. 10.1073/pnas.2117270119

50. Bamford, N.S., Wightman, R.M., and Sulzer, D. (2018). Dopamine’s Effects on Corticostriatal Synapses during Reward-Based Behaviors. Neuron 97, 494–510. 10.1016/j.neuron.2018.01.006.

51. Han, D.H., Park, P., Choi, D.I., Bliss, T.V.P., and Kaang, B.K. (2022). The essence of the engram: Cellular or synaptic? Seminars in cell & developmental biology 125, 122–135. 10.1016/j.semcdb.2021.05.033.

52. Gipson, C.D., Kupchik, Y.M., Shen, H., Reissner, K.J., Thomas, C.A., and Kalivas, P.W. (2013). Relapse induced by cues predicting cocaine depends on rapid, transient synaptic potentiation. Neuron 77, 867–872. 10.1016/j.neuron.2013.01.005.

53. Cruz, F.C., Babin, K.R., Leao, R.M., Goldart, E.M., Bossert, J.M., Shaham, Y., and Hope, B.T. (2014). Role of Nucleus Accumbens Shell Neuronal Ensembles in Context-Induced Reinstatement of Cocaine-Seeking. J. Neurosci. 34, 7437–7446. 10.1523/Jneurosci.0238-14.2014.

54. Caprioli, D., Venniro, M., Zhang, M., Bossert, J.M., Warren, B.L., Hope, B.T., and Shaham, Y. (2017). Role of Dorsomedial Striatum Neuronal Ensembles in Incubation of Methamphetamine Craving after Voluntary Abstinence. J Neurosci 37, 1014–1027. 10.1523/JNEUROSCI.3091-16.2016.

55. Pribiag, H., Shin, S., Wang, E.H., Sun, F., Datta, P., Okamoto, A., Guss, H., Jain, A., Wang, X.Y., De Freitas, B., et al. (2021). Ventral pallidum DRD3 potentiates a pallido-habenular circuit driving accumbal dopamine release and cocaine seeking. Neuron 109, 2165–2182 e2110. 10.1016/j.neuron.2021.05.002.

56. Bobadilla, A.C., Heinsbroek, J.A., Gipson, C.D., Griffin, W.C., Fowler, C.D., Kenny, P.J., and Kalivas, P.W. (2017). Corticostriatal plasticity, neuronal ensembles, and regulation of drug-seeking behavior. Brain Research in Addiction 235, 93–112. 10.1016/bs.pbr.2017.07.013.

57. Hong, I., Song, B., Lee, S., Kim, J., Kim, J., and Choi, S. (2009). Extinction of cued fear memory involves a distinct form of depotentiation at cortical input synapses onto the lateral amygdala. Eur J Neurosci 30, 2089–2099. 10.1111/j.1460-9568.2009.07004.x.

58. Lee, J.H., Kim, W.B., Park, E.H., and Cho, J.H. (2022). Neocortical synaptic engrams for remote contextual memories. Nat Neurosci. 10.1038/s41593-022-01223-1.

59. Kim, J., Lee, S., Park, K., Hong, I., Song, B., Son, G., Park, H., Kim, W.R., Park, E., Choe, H.K., et al. (2007). Amygdala depotentiation and fear extinction. Proc Natl Acad Sci U S A 104, 20955–20960. 10.1073/pnas.0710548105.

60. Pettibone, J.R., Yu, J.Y., Derman, R.C., Faust, T.W., Hughes, E.D., Filipiak, W.E., Saunders, T.L., Ferrario, C.R., and Berke, J.D. (2019). Knock-In Rat Lines with Cre Recombinase at the Dopamine D1 and Adenosine 2a Receptor Loci. Eneuro 6. Artn 0163-19.2019 10.1523/Eneuro.0163-19.2019.

61. Wei, X., Ma, T., Cheng, Y., Huang, C.C.Y., Wang, X., Lu, J., and Wang, J. (2018). Dopamine D1 or D2 receptor-expressing neurons in the central nervous system. Addiction biology 23, 569–584. 10.1111/adb.12512.

62. Lu, J., Cheng, Y., Wang, X., Woodson, K., Kemper, C., Disney, E., and Wang, J. (2019). Alcohol intake enhances glutamatergic transmission from D2 receptor-expressing afferents onto D1 receptor-expressing medium spiny neurons in the dorsomedial striatum. Neuropsychopharmacology 44, 1123–1131. 10.1038/s41386-019-0332-9.

63. Cheng, Y., Xie, X., Lu, J., Gangal, H., Wang, W., Melo, S., Wang, X., Jerger, J., Woodson, K., Garr, E., et al. (2021). Optogenetic induction of orbitostriatal long-term potentiation in the dorsomedial striatum elicits a persistent reduction of alcohol-seeking behavior in rats. Neuropharmacology 191, 108560. 10.1016/j.neuropharm.2021.108560.

64. Huang, C.C.Y., Ma, T., Roltsch Hellard, E.A., Wang, X., Selvamani, A., Lu, J., Sohrabji, F., and Wang, J. (2017). Stroke triggers nigrostriatal plasticity and increases alcohol consumption in rats. Sci Rep 7, 2501. 10.1038/s41598-017-02714-z.

65. Visser, E., Matos, M.R., van der Loo, R.J., Marchant, N.J., de Vries, T.J., Smit, A.B., and van den Oever, M.C. (2020). A persistent alcohol cue memory trace drives relapse to alcohol seeking after prolonged abstinence. Sci Adv 6, eaax7060. 10.1126/sciadv.aax7060.

66. Xie, X., Chen, R., Wang, X., Smith, L., and Wang, J. (2023). Activity-dependent labeling and manipulation of fentanyl-recruited striatal ensembles using ArcTRAP approach. STAR Protoc 4, 102369. 10.1016/j.xpro.2023.102369.

67. Franklin, K.B.J., and Paxinos, G. (2008). The mouse brain in stereotaxic coordinates, 3rd Edition (Boston : Elsevier/Academic Press).

68. Gangal, H., Xie, X., Huang, Z., Cheng, Y., Wang, X., Lu, J., Zhuang, X., Essoh, A., Huang, Y., Chen, R., et al. (2023). Drug reinforcement impairs cognitive flexibility by inhibiting striatal cholinergic neurons. Nature communications 14, 3886. 10.1038/s41467-023-39623-x.

